# Automated satellite remote sensing of giant kelp at the Falkland Islands (Islas Malvinas)

**DOI:** 10.1101/2021.09.14.460404

**Authors:** Henry F. Houskeeper, Isaac S. Rosenthal, Katherine C. Cavanaugh, Camille Pawlak, Laura Trouille, Jarrett E.K. Byrnes, Tom W. Bell, Kyle C. Cavanaugh

## Abstract

Giant kelp populations support productive and diverse coastal ecosystems in both hemispheres at temperate and subpolar latitudes but are vulnerable to changing climate conditions as well as direct human impacts. Observations of giant kelp forests are spatially and temporally patchy, with disproportionate coverage in the northern hemisphere, despite the size and comparable density of southern hemisphere kelp forests. Satellite imagery enables the mapping of existing and historical giant kelp populations in understudied regions, but automating the detection of giant kelp in large satellite datasets requires approaches that are robust to the optical complexity of the shallow, nearshore environment. We present and compare two approaches for automating the detection of giant kelp in satellite datasets: one based on crowd sourcing of satellite imagery classifications and another based on a decision tree paired with a spectral unmixing algorithm (automated using Google Earth Engine). Both approaches are applied to satellite imagery (Landsat) of the Falkland Islands or Islas Malvinas (FLK), an archipelago in the southern Atlantic Ocean that supports expansive giant kelp ecosystems. The performance of each method is evaluated by comparing the automated classifications with a subset of expert-annotated imagery cumulatively spanning over 2,700km of coastline. Using the remote sensing approaches evaluated herein, we present the first continuous timeseries of giant kelp observations in the FLK region using Landsat imagery spanning over three decades. We do not detect evidence of long-term change in the FLK region, although we observe a recent decline in total canopy area from 2017-2021. Using a nitrate model based on nearby ocean state measurements obtained from ships and incorporating satellite sea surface temperature products, we find that the area of giant kelp forests in the FLK region is positively correlated with the nitrate content observed during the prior year. Our results indicate that giant kelp classifications using citizen science are approximately consistent with classifications based on a state-of-the-art automated spectral approach. Despite differences in accuracy and sensitivity, both approaches find high interannual variability that impedes the detection of potential long-term changes in giant kelp canopy area, although recent canopy area declines are notable and should continue to be monitored carefully.

## Introduction

Kelp forests make up an important coastal habitat that provides productive and dynamic biotic structure, supports diverse marine ecosystems and fisheries, and supplies resources to coastal communities [1, 2]. These coastal ecosystems are extensive across temperate to sub-polar latitudes of both hemispheres. The largest and most widely distributed canopy-forming kelp is giant kelp, *Macrocystis pyrifera*. Giant kelp extends along the west coast of North America and throughout much of the southern hemisphere, including along the South American coastlines of Peru, Chile, and Argentina, the southern margins of the African and Australian continents, and various island chains situated at southern temperate and sub-antarctic latitudes [3, 4].

The global distribution of giant kelp is regulated by biotic and abiotic factors, including: the depth and quality of the benthic substrate; the temperature, clarity, and nutrient content of the water; the magnitude and direction of wave energy; the size and connectivity of kelp populations; and the abundance of grazers, notably sea urchins [5–7]. Kelp population responses to specific environmental drivers are nonlinear, and the relative importance of biotic and abiotic controls may vary seasonally and regionally, even across small distances due in part to complex variations in coastline structure and exposure [8, 9]. The sensitivity of kelp to environmental drivers leads to high variability in kelp forest abundance across a variety of space and time scales [6]. This sensitivity also makes kelp ecosystems especially vulnerable to changes in environmental conditions. As a result, regular monitoring is required to detect potential changes in the distribution of giant kelp corresponding with environmental changes, and long-term data sets are needed to separate trends from natural background variability [10].

### Global and regional changes in kelp forest distribution

Giant kelp forest timeseries observations are geographically patchy, with more research focusing on the northern hemisphere (especially southern Californian waters) [3]. Two recent surveys of kelp forests at the South Atlantic Tristan da Cunha Islands and at the southern tip of Chile reported highly dense kelp forests with ecosystem qualities comparable to those of the well-studied Marine Protected Areas (MPAs) of southern California [11, 12]. The forests of the South Atlantic region comprise a significant fraction of the global giant kelp distribution; a subset of key southern hemisphere ecosystems that included Tierra del Fuego, the Falkland Islands (Islas Malvinas; FLK), and the South Georgia Islands was recently estimated to comprise approximately 41% of the global coverage of giant kelp [4]. Despite the significant coverage of giant kelp in the South Atlantic, there are insufficient observations from many southern hemisphere regions – including FLK – to determine whether South Atlantic forests have exhibited long term trends related to climate change [13, 14].

The giant kelp forests of the FLK region, which consists of an archipelago situated approximately 600km east of Tierra del Fuego, are believed to be one of the largest remaining, relatively undisturbed kelp forest ecosystems on the planet [15]. The economic value of these forests was recently estimated at £2.69 billion based on carbon sequestration, coastal protection, and other ecosystem services [16]. Regions like FLK – which do not support large human population centers, have not recently undergone large changes in land-use practices, and have relatively low levels of local pollution or resource extraction – are useful for examining the impacts of climate change because the effects of local environmental stressors are reduced. Despite relatively few modern observations, giant kelp canopies in FLK were among the first kelp forests to be documented by naturalists during the 19^th^ and early 20^th^ centuries [15] and were the subject of a series of physiological and ecological studies during the latter portion of the 20^th^ century [17–20]. An update on the status of giant kelp forests within the FLK region – based on continuous and spatially resolved kelp canopy observations – is timely given uncertainty in the resilience of remote South Atlantic kelp populations and would enable better assessment of long-term variability as well as key regional environmental controls for this valuable region.

### Advances in observing remote kelp ecosystems

Detecting robust long-term change in kelp abundance requires persistent, regular monitoring of kelp forest ecosystems, which is only feasible on small spatial scales when using local surveys, e.g., within a Southern California ecological reserve [21]. For less well-characterized regions, comparison of present conditions with those documented in historical surveys allows for useful point comparisons but cannot resolve interannual variability, the magnitude of which can often overshadow long-term changes. Satellite data improve the temporal and spatial coverage of observations of kelp forest ecosystems [22] and enable historical reconstruction of kelp canopy area spanning multiple decades [10]. Satellite remote sensing has been used to assess environmental drivers [9], understand the importance of decadal-scale processes in the regulation of kelp forests [10], and detect long-term trends [23].

As the proliferation of ocean-viewing, high-resolution satellite imagers have greatly increased the volume of data relevant to observing kelp canopies, improvements in image processing and automation are needed to apply satellite observations across broader geographic areas. Recent technological developments have eased image processing requirements, e.g., the Google Earth Engine (GEE) platform facilitates large-scale data analysis through cloud computing [24]. For example, a global map of kelp forest canopy was recently generated by processing curated, satellite imagery within the GEE cloud platform [4].

Advances in processing satellite datasets have also resulted from novel approaches based on crowd sourcing, in which citizen scientists can accomplish simple, repetitive tasks that would otherwise be too time-consuming for a small research team to complete [25], e.g., manually annotating large sets of satellite imagery. These citizen science annotations can also be used as training and validation data for automated machine learning classification approaches. This concept of including public effort in scientific research also connects the general public with teams of scientists, thereby facilitating general science literacy and engagement [26, 27]. In the field of environmental science, crowd sourcing has been utilized for decades, but has also faced concerns regarding the quality and consistency of crowd-sourced data products [28]. These concerns can be mitigated to some extent through optimizing the user-interface, cleaning the data using post-processing tools, expanding the platform’s accessibility, and improving communication between scientists and participants [29].

Kelp canopy floating at the ocean’s surface is often clearly distinguishable to an unskilled human observer, especially for pseudo- or false-color imagery in which the near-infrared (NIR) signal is emphasized. Crowd-sourced kelp annotations of satellite imagery were generated by the Floating Forests (FF) project (floatingforests.org) beginning in 2014 through the Zooniverse citizen science web portal (zooniverse.org). Using validation data from southern California, the FF kelp canopy data products have recently been shown to be comparable in quality to manual, expert annotations when filtered by consensus (a minimum ratio of positive annotations for an individual pixel) [30]. This project has expanded geographically, and as of the writing of this paper, the kelp canopy annotations completed through the FF project span the coastal regions of California, the western edge of Tasmania, and the FLK archipelago. In FLK, over 433 images have presently been classified by citizen scientists.

In this paper, we present the first continuous timeseries of giant kelp observations in the FLK region using over three decades of Landsat imagery. We derived canopy area estimates from imagery based on multiple classification approaches, including citizen science classifications obtained through the FF project, as well as a fully-automated, two-part analytical approach, which pairs a binary decision tree with a spectral mixing model [10]. The addition of the spectral mixing model allows for assessment of partially covered pixels and has been shown to improve detection for canopies with a smaller surface footprint [31]. We evaluated the automated and crowd-sourced kelp classifications using an expert-annotated imagery subset, and we tested for long-term changes in canopy distribution using multisensor timeseries of the FLK region. Finally, we assessed potential environmental drivers of variability in FLK giant kelp populations using independent data products of sea surface temperature (SST) and nitrate-temperature relationships approximated from nearby oceanographic surveys.

## Materials and methods

### Site description

The FLK region constitutes an archipelago of over 700 south Atlantic islands. The region’s two largest islands, West and East Falkland, are separated by Falkland Sound, which is oriented approximately southwest to northeast along its longest axis. The archipelago is situated in a zone of high wave energy, with maximum wave heights occurring in the austral winter months and approaching from the west [32]. Coastline morphology is strongly affected by wind and wave exposure, with high rates of erosion along the western, windward coastline of many islands, and sediment deposits or formation of dunes more common along the eastern, leeward coastlines [33].

The ocean state of the FLK region is strongly influenced by the flow of the Falkland Current (FC), which supplies cold (~ 7°C) water originating in the Antarctic Circumpolar Current (ACC), shown in Fig 1. To the north of FLK, between approximately 30° and 50°, the FC forms a confluence with warmer (~ 19°C) and more saline waters of the Brazil Current (BC) [32]. Tidal cycles in FLK are semi-diurnal, with ranges of approximately 1m [32], and the region’s subtidal zone supports two species of sea urchins that graze kelp [34].

**Fig 1.**
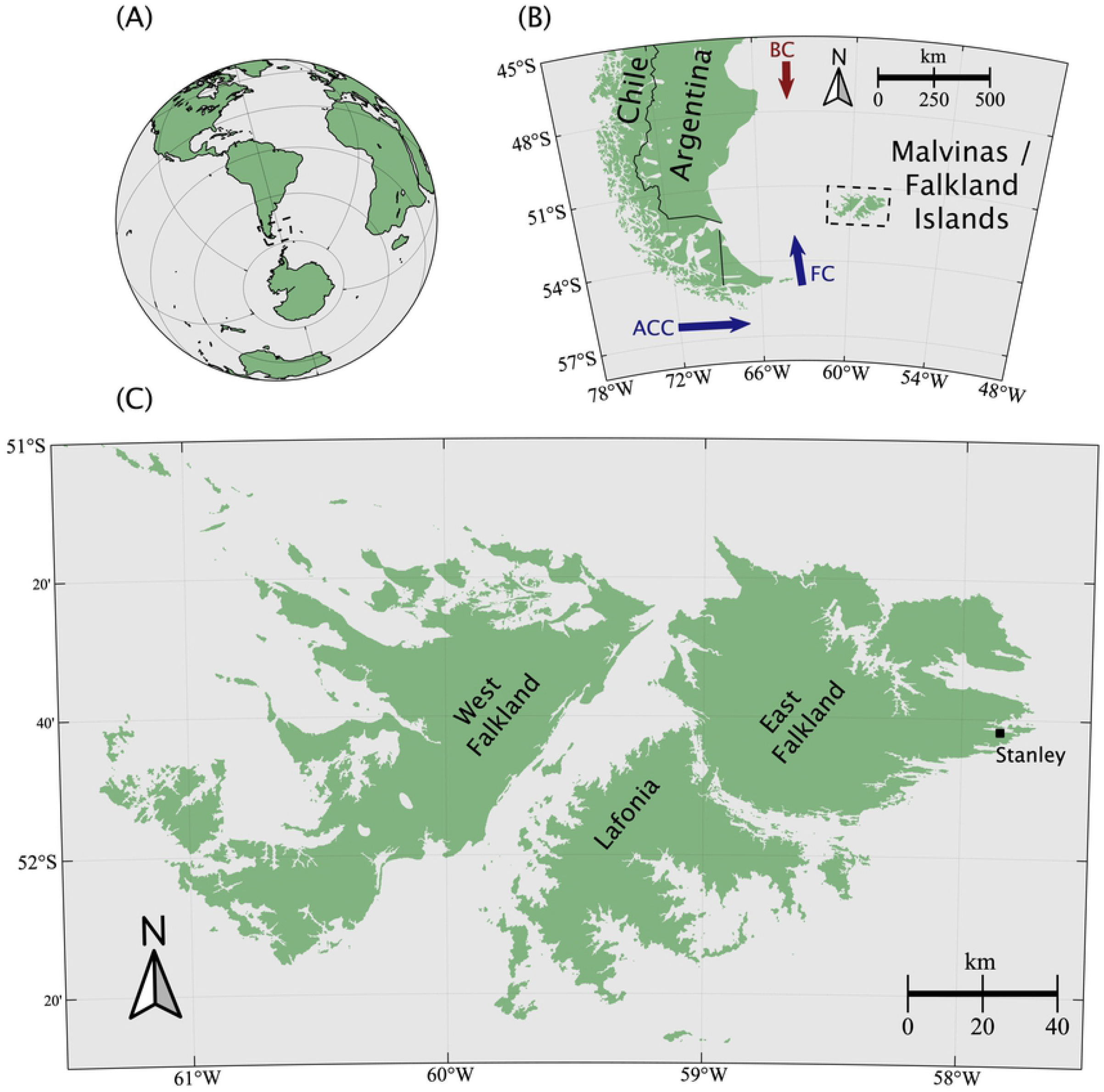
Map of Malvinas or Falkland Islands. The location and coastline of the Falkland Islands (Islas Malvinas) are shown at increasing scale within panels (A), (B), and (C). Within panel (B), the generalized location and direction of the Antarctic Circumpolar Current (ACC) and the Falkland Current (FC) are indicated in blue, and the Brazil Current (BC) is indicated in red.

Giant kelp is the dominant canopy-forming kelp within the FLK region, with large beds surrounding the island archipelago [17]. Other canopy-forming genera in the FLK region are primarily found in very shallow habitats or as a subsurface canopy interspersed within or on the edges of giant kelp beds [15, 34]. For example, *Lessonia* spp. frequently forms a sub-canopy that rarely reaches the surface except in very shallow waters, and the southern bull kelp *Durvillaea antarctica* is generally restricted to rocky shore margins or tidally submerged reefs. Giant kelp fronds in FLK exhibit rapid turnover, with a maximum estimated frond lifespan of one year [18], and individuals in nearshore beds have been found to experience nitrogen limitation in summer months [17]. Recent genetic analyses have revealed that giant kelp populations in the FLK region share commonalities with populations of central Chile, suggesting that giant kelp arrived in FLK prior to a recolonization event of southern Chile [35].

### Satellite mapping of kelp dynamics

We obtained satellite remote sensing observations of the FLK region using available Landsat 5, 7, and 8 imagery. We extracted kelp canopy area from the Landsat imagery using a citizen-science approach [30], as well as the current state-of-the-art automated method [10]. The two different approaches provided redundancy and also allowed us to compare each method using manual, expert classifications as a validation dataset. Based on the validation results presented in this study, we applied the best performing method to examine variability in kelp abundance across FLK from 1985 - 2021. The satellite imagery and giant kelp data products based on expert, automated, and citizen science classifications are described in more detail below.

### Expert kelp classification of Landsat imagery

Atmospherically corrected surface reflectance observations of the FLK region were obtained from the Landsat satellites via the United States Geological Survey (USGS) Earth Explorer portal (`earthexplorer.usgs.gov`). Scenes were downloaded from Landsat Collection 1 for the Thematic Mapper (TM), Enhanced Thematic Mapper Plus (ETM+), and Operational Land Imager (OLI) sensors, which were carried aboard Landsat platforms 5, 7, and 8, respectively. Kelp-containing pixels (30m spatial resolution) were manually classified by a single, skilled technician using pseudo-color (NIR/red/green) imagery for a subset of eight Landsat scenes, cumulatively spanning over 2,700km of coastline. The manually annotated scenes were primarily cloud-free and with similar representation from each sensor (2 TM, 3 ETM+, and 3 OLI), but were otherwise chosen to be representative of the broader dataset, with dates spanning: a) the austral spring, summer, and fall (winter observations were avoided due to solar zenith angle); b) the majority (1999 – 2018) of the continuous timeseries range (1997 – 2021); and c) a 1.6 σ range in the El Niño Southern Oscillation (ENSO), using the Multivariate ENSO Index (MEI). Manual classifications were compared to automated and citizen science classifications by evaluating the number of pixels identified as kelp for the same Landsat scenes, with pixels partitioned into bins corresponding to the nearest 1km FLK coastline segment.

### Automated kelp classifications based on a spectral mixture analysis

Atmospherically corrected surface reflectance products (tier 1) were obtained for Landsat sensors TM, ETM+, and OLI from the GEE public data catalogue. Within the GEE code editor (code.earthengine.google.com), scenes were masked for land and clouds, and kelp-containing pixels were identified by following a binary classification scheme described in Bell et al., 2020 [10]. The fractional kelp coverage within each kelp-containing pixel was then estimated based on Bell et al., 2020 [10] using Multiple Endmember Spectral Mixture Analysis (MESMA) – a technique for estimating fractional contributions of pure spectral endmembers (e.g., water, glint, or kelp) from individual bulk spectra based on a linear mixing model [36]. The water end-members were derived from the imagery, while a single kelp end-member was chosen *a priori*.

The data products were then post-processed in Matlab (R2020a) by masking pixels in locations that were characterized as follows: located less than 120m or greater than 4.5km from the coastline, defined by [37]; containing kelp canopy in less than 2% of the timeseries; containing fractional kelp coverage assignments above 300%; indicating strong negative (< –0.25) correlation to tidal cycles estimated from sea surface pressure observations at Port Stanley [38], which were accessed through the University of Hawaii Sea Level Center (`uhslc.soest.hawaii.edu`); and/or indicating low (<0.2) correlation with nearby (within 5km) patches. Filtering the automated classifications beyond the post-processing criteria outlined above was not performed, since the objective of our paper is in part to evaluate the accuracy of an automated remote approach. The fractional cover datasets were converted to binary timeseries (i.e., presence/absence) – based on a fractional coverage threshold of 13% – in order to facilitate comparisons with the expert and citizen science classifications. Previous work has shown that spectral response functions have improved intersensor consistency for non-binary, percent-coverage data products [10]. However, spectral response functions were not applied here because inter-sensor differences were not detected for DTM products, which were binary, i.e., presence-absence rather than fractional. This dataset of binary kelp canopy classifications was re-sampled at seasonal or annual (July - June) intervals using the mean and is hereafter referred to as DTM. The number of observation days in the annual DTM dataset is shown in Supplemental Sect. S1.

Another automated approach based on the difference between NIR and red satellite reflectance observations was also processed within the GEE code editor using open-access code (github.com/BiogeoscienceslabOxford/kelp_forests). Atmospherically corrected surface reflectance products were obtained from the GEE public data repository for the MultiSpectral Imager (MSI) aboard the European Space Agency satellite Sentinel-2. Differences between NIR and red reflectance products were converted to binary presence/absence kelp data products based on predetermined thresholds [4]. The open-access code was adjusted to produce annual composites, and this dataset is hereafter referred to as KD. The KD dataset is included for visual comparisons but is not included in the validation analysis, since the algorithm was not applicable to the Landsat imagery on which the manual, expert classifications were performed.

### Citizen science kelp classifications

Crowd-sourced satellite kelp annotations were obtained for TM, ETM+, and OLI Landsat imagery (USGS tier 1) through the FF project (floatingforests.org), in which scenes were spatially subset into smaller tiles (≈ 2.25km^2^), and color-stretched tile images (red/green/blue) were annotated by citizen scientists through the Zooniverse citizen science web portal (zooniverse.org), shown in Fig. 2. Each tile was viewed by 15 unique, unskilled observers during the period 2014 to 2020, with annotations recorded as shape files. Annotated tiles were combined as rasterized (10m resolution) data composites containing the summed annotations for each pixel within each spatially subset tile.

**Fig 2.**
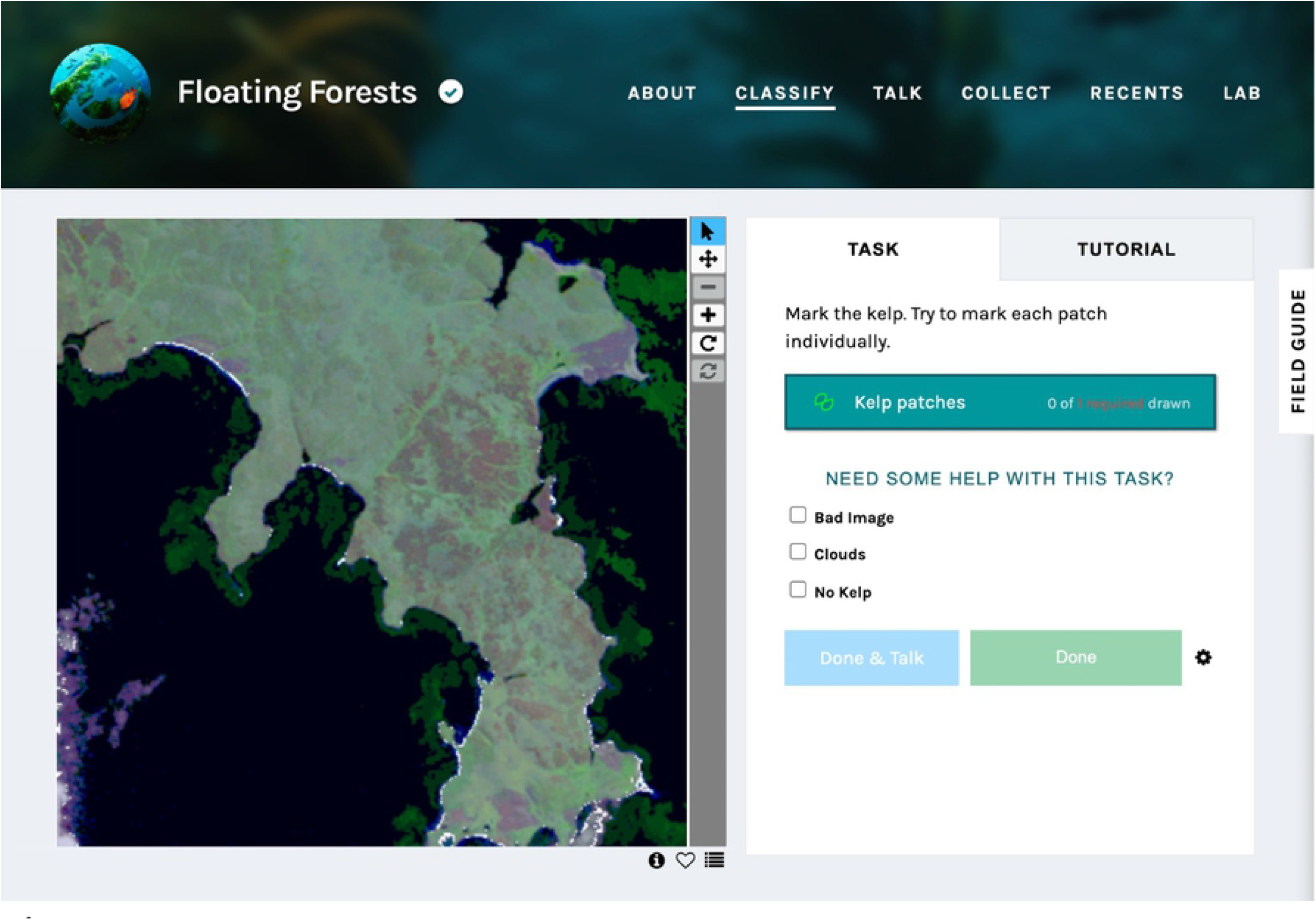
Zooniverse citizen science web portal. The Zooniverse interface in which citizen scientists indicate kelp canopy locations for the Floating Forests project. A representative tile viewing a portion of East Falkland is shown.

Tiles were then post-processed using Matlab by rasterizing the shapefiles and then converting to binary using a consensus classification threshold of 8. The consensus threshold was selected based on visual inspection of scenes as well as optimization of the Matthew’s Correlation Coefficient – a metric that is relatively insensitive to class imbalance [39] – in comparison to expert-annotated imagery. Binary annotations that were on land or that were greater than 4.5km from the nearest coastline were discarded. In general, oceanic consensus classifications were included without additional filtering, since our objective is in part to evaluate the accuracy of the citizen science data products. However, one tile was determined to contain primarily drifting kelp or marine debris and was manually removed from the dataset during post-processing. Binary rasters were re-sampled to a 30m grid and to seasonal or annual intervals using the mean. Seasonal or annual composites with insufficient data (less than 25%) were removed from the time series analysis. This dataset of binary kelp canopy classifications based on the Floating Forest data set and the consensus threshold of 8 is hereafter referred to as FF8. The number of observations days in the annual FF8 dataset is shown in Supplemental Sect. S1.

### Ocean state estimates

Monthly, 9km composites of sea surface temperature (SST) were obtained from the NASA Ocean Color portal (accessed through `oceancolor.gsfc.nasa.gov`) using observations from the Moderate Resolution Imaging Spectroradiometer (MODIS) aboard NASA’s *Aqua* satellite. Observations spanning 2002 to 2021 were averaged within 62° to 57° W and 50° to 53° S. Oceanographic observations of nitrate and temperature were obtained for the same region from the World Ocean Atlas – accessed through the National Centers for Environmental Information (`ncei.noaa.gov`). A synthetic nitrate proxy was modeled as a function of temperature using linear regression, with estimated uncertainty (RMSE) of 4.6 *μ*mol/kg, or 13.1% of the range in nitrate measured *in situ* (Fig. S2).

## Results

We compared automated and citizen science approaches for estimating giant kelp canopy coverage and evaluated the performance of each method for the FLK region based on validation against expert annotations. We developed kelp canopy time series for the FLK region and tested for associations between giant kelp area and environmental parameters or climate indices using the best performing approach. Finally, we investigated regional relationships by aggregating kelp canopy data products to the nearest 1km coastline segments.

### Evaluation of kelp canopy data products

Kelp classifications from the automated approach (DTM) matched the expert manual classifications better than the citizen science consensus (FF8). Fig 3 shows similar kelp canopy distributions estimated using DTM data products from separate Landsat sensors OLI and ETM+, shown in panels C and D, respectively. The FF8 citizen science classifications are shown in panel E and indicate decreased spatial granularity as well as decreased sensitivity to small beds. KD data products generated from Sentinel-2 (MSI) imagery are shown in panel F and indicate slightly less sensitivity than the DTM approach, although the DTM and KD methods are approximately similar in shape and in total coverage.

**Fig 3.**
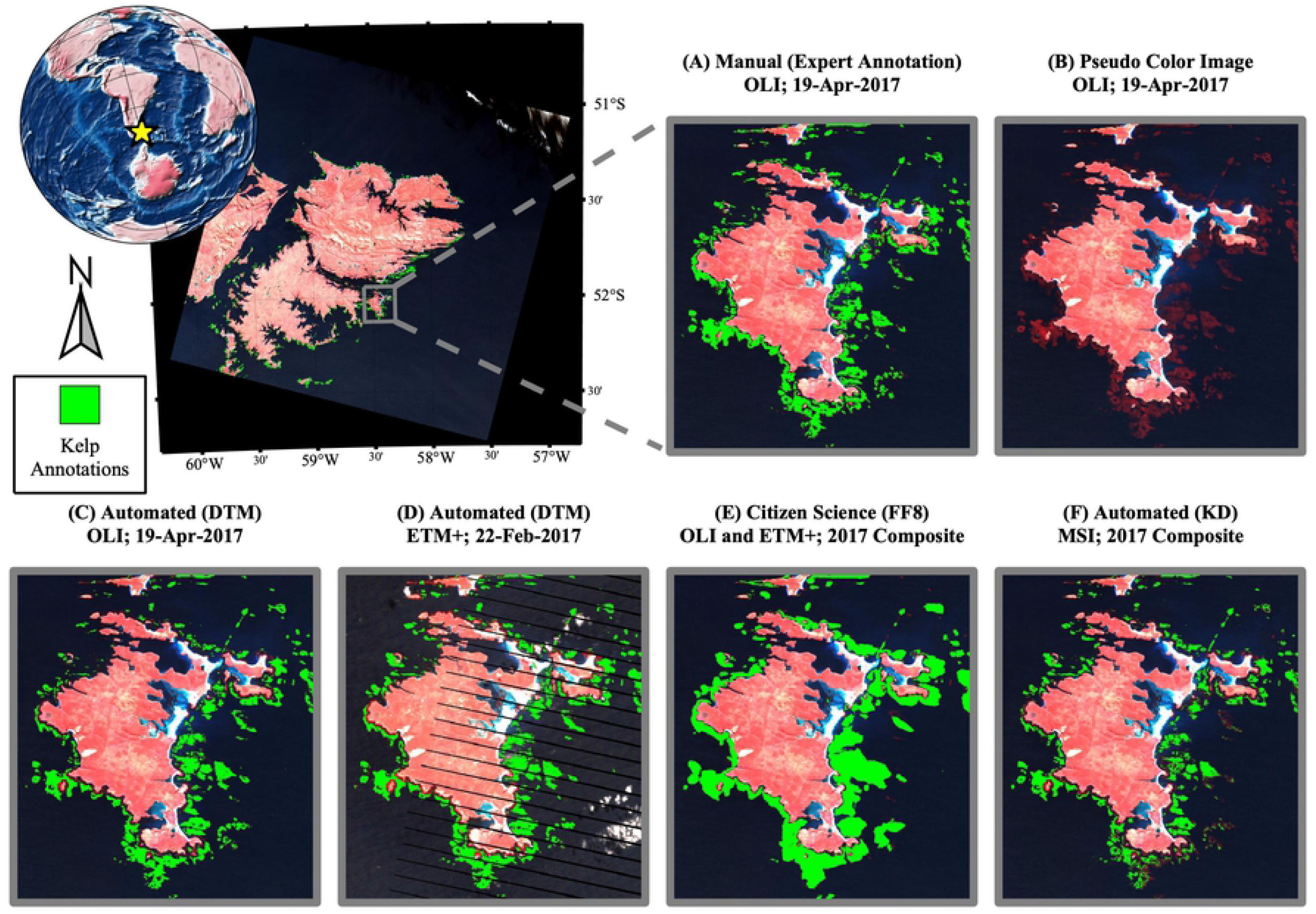
Comparison of kelp classification methods. A pseudo-color scene of East Falkland is shown in the upper left panel with expert kelp classifications indicated in green. The figure inlays show various automated and manual methods, as well as non-annotated imagery, for a smaller spatial region, as follows: (A) manual classifications performed on a single OLI scene by an expert technician; (B) pseudo-color (NIR/red/green) image; (C) DTM automated classifications of a single OLI scene; (D) DTM automated classifications of a single ETM+ scene; (E) FF8 consensus classifications of OLI imagery during 2017; and (F) KD automated classifications of MSI imagery during 2017.

The automated DTM and KD approaches were based on spectral analysis and were not retained for near-shore pixels (i.e., within 120m or approximately 4 pixel-widths from the coastline), where bottom reflectance or the additions of organic or inorganic particles (e.g., through resuspension or terrestrial inputs) challenge atmospheric correction [40] and elevate signals in the NIR domain [41]. In some instances, nearshore remote sensing challenges are anticipated to be less problematic when spatial information is also considered (e.g., the trajectory of a runoff plume), and so the FF8 annotations were retained within the nearshore zone. Continued research is needed in order to better understand the reliability of spectral methods like DTM and KD for measuring nearshore canopy. The FLK region, in general, contains relatively wide giant kelp canopies that extend away from the shore [15], and improvements for the nearshore zone are particularly important in regions where kelp predominantly forms fringing canopies closer to the shoreline.

Quantitative comparison of the DTM and FF8 methods was performed by aggregating kelp-containing pixels to their nearest 1km coastline segment and comparing the number of classifications within each segment with the corresponding number of expert classifications. Validation results are presented in Fig 4, with the DTM approach shown using the TM, ETM+, and OLI sensors in panels A, B, and C, respectively. Panel D shows the FF8 approach using the TM and OLI sensors overlaid, due to the lower total number of available matchups. No ETM+ matchups were available for coincident FF8 data products and expert-annotated imagery.

**Fig 4.**
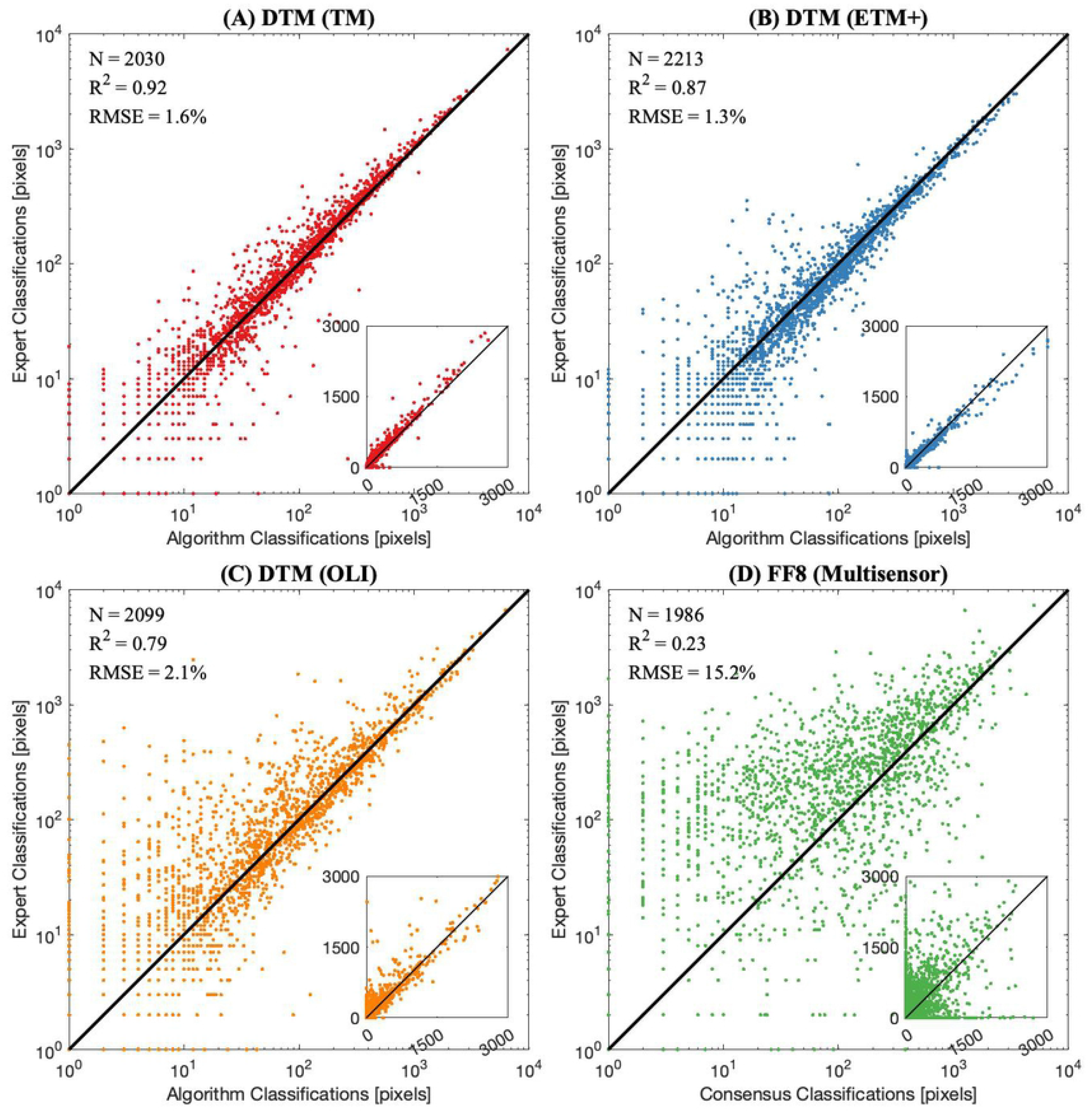
Validation of automated and citizen science kelp classifications. Validation scatterplots for expert (vertical) and automated or citizen science consensus (horizontal) classifications of pixels with kelp present per 1km of coastline, as follows: (A) validation of TM matchups using DTM; (B) validation of ETM+ matchups using DTM; (C) validation of OLI matchups using DTM; and (D) validation of OLI and TM matchups using FF8.

Across all sensors, the DTM approach produced the highest R^2^ and lowest RMSE, expressed as a percentage of the range in the expert-annotated coastline aggregates, compared to the FF8 data products. For consistency, the challenging nearshore pixels that were masked for the DTM data products were also removed in the validation of the FF8 aggregates. Biases of the FF8 data products within individual matchup scenes were greater in magnitude than those of the DTM data products, but occurred in both directions across scenes. Optimizing sensitivity and specificity for the citizen science classifications is possible through adjusting the consensus threshold. We found that a consensus threshold of 8 was reasonable within the FLK region, but another study recently found that a consensus threshold of 4 (i.e., more sensitivity) was more appropriate based on annotated imagery of Southern California waters [30]. Different optimization results may be due in part to the increased width and offshore extent of the giant kelp beds in FLK, particularly along the eastern coastline. The different optimization results may also be due to differences in image processing, because the earlier imagery available to Rosenthal et al., (2018) [30] was not atmospherically corrected.

### Interannual variability in kelp canopy extent (1985-2021)

Timeseries of total FLK kelp canopy coverage derived from the FF8 and DTM approaches indicate that the spectral and citizen science approaches show similar temporal patterns and trends. Annual composites of total kelp canopy area are shown in Fig. 5, with DTM data products shown for individual sensors TM, ETM+, and OLI in red, blue, and orange, respectively, and for a multisensor composite in gray. FF8 data products are shown as an annual, multisensor composite in green due to the lower amount of available observations. Datasets for individual sensors were combined as multisensor composites based on a weighted mean that incorporated the number of observations available from each sensor within a single year (see Supplemental Table S1). The earliest DTM observations are from 1985, but coverage during this year only includes the western portion of FLK.

**Fig 5.**
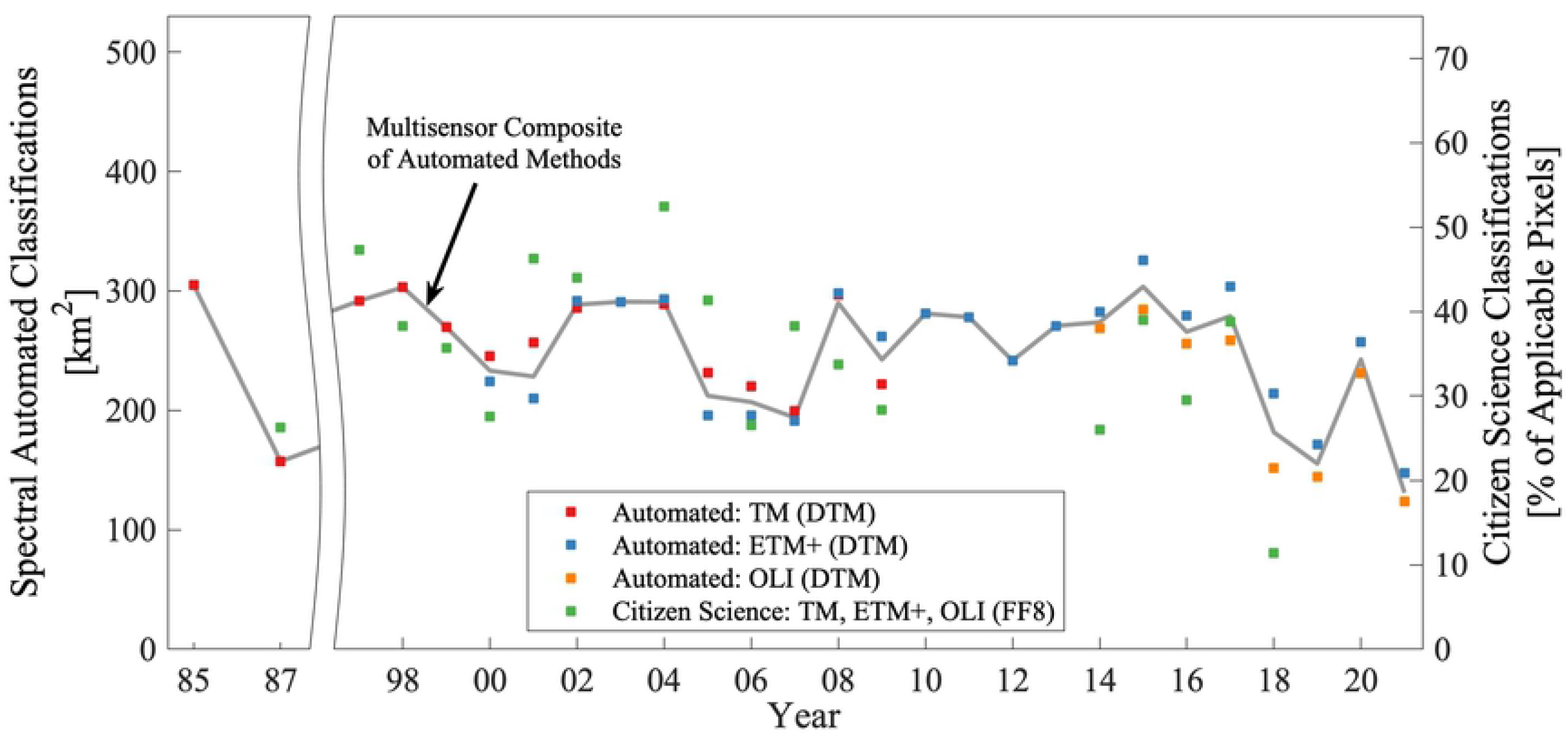
Timeseries of kelp canopy extent for the combined FLK region. Annual mean total canopy area within the FLK region is shown for DTM data products using TM (red), ETM+ (blue), and OLI (yellow) individual sensors, as well as a multisensor composite (gray line). FF8 data products are scaled to a separate y-axis based on the percentage of applicable pixels observed in an individual year, due to spatial patchiness in the FF8 coverage through time, and are presented as a multisensor composite (green). DTM observations in 1985 only include the western portion of FLK. Yearly intervals correspond to July of the preceding calendar year through June of the listed calendar year.

We did not detect significant long-term linear trends in kelp canopy using either the DTM data products spanning 1985-2021 (P = 0.28) or the FF8 data products spanning 1987-2018 (P = 0.09). Recent declines in kelp coverage are apparent for the full FLK region during 2017-2020, but the magnitude of this recent decline is similar to the range observed in the DTM timeseries between 1985 and 1987. We also tested for regional kelp canopy trends by evaluating timeseries of DTM data products aggregated to their nearest 1km coastline segments. Considering timeseries results from each 1km segment by coastline orientation and position, we did not detect cohesive spatial structure for trends in kelp canopy (Fig. S3). Approximately 3.9% and 8.8% of the 1km coastline partitions indicated significant (P < 0.01) positive and negative linear trends, respectively.

We also evaluated potential effects of basin-scale climate indices by testing for lag correlation with annual and seasonal DTM data products. Limiting the lag in the climate index to one year (i.e., because of the rapid turnover expected for giant kelp populations), no significant correlation was detected with MEI or other indices.

Using a synthetic nitrate model described in Supplemental Section S2, we found that annual composites of kelp canopy area were positively correlated to nitrate concentration when nitrate was lagged by one year, with r = 0. 65 (P < 0. 01) for the combined FLK region. Correlations derived between canopy area and synthetic nitrate within individual, 1km coastline subsets indicated that positive associations between kelp and lagged nitrate were consistent across most FLK coastline segments, shown in Fig. 6. Similar results were obtained for temperature because of the linear model used to estimate synthetic nitrate, but only nitrate is presented because temperatures in FLK are not anticipated to reach levels associated with thermal stress in giant kelp.

**Fig 6.**
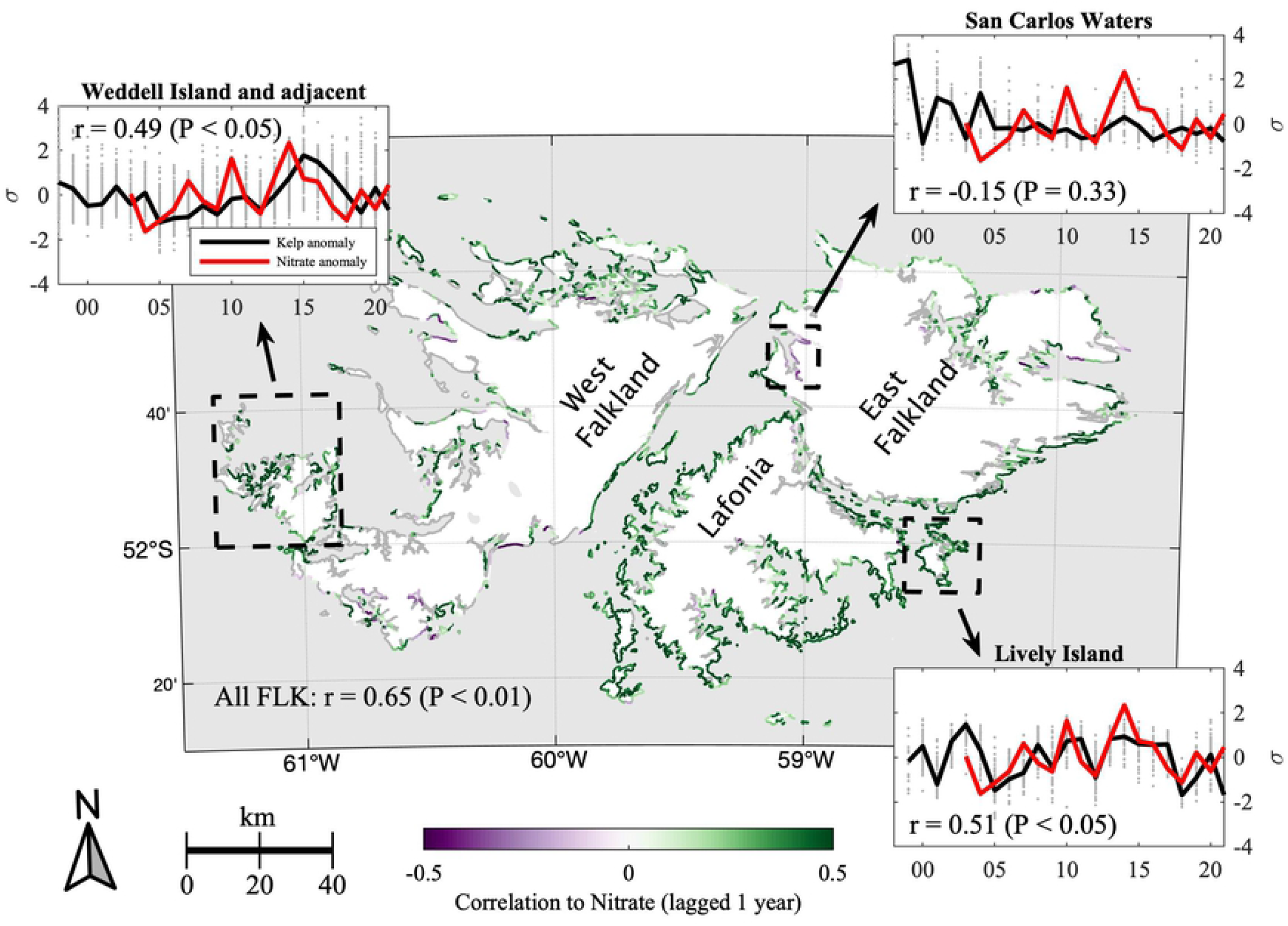
Spatial variability in correlation between standardized canopy extent and modeled nitrate concentration. Correlation coefficients for modeled nitrate concentrations and DTM standardized canopy extents are shown in purple and green for 1km coastline subsets. If P ≥ 0.01, coefficients are instead indicated in gray. Timeseries of select coastline subsets are shown in the figure inlays, with standardized kelp canopy extent indicated in gray for individual 1km coastlines and in black for the combined sub-region, and with modeled annual nitrate shown in red. Years are indicated on the x-axis. The sub-regions included in the figure inlays are indicated with a dashed black line, and correspond as follows: Weddell Island and adjacent islands (upper left); San Carlos waters (upper right); and Lively Island (lower right).

Areas where nitrate correlations were either negative or were not found to be significant (shaded purple or gray, respectively) were typically in regions with very low or ephemeral kelp canopy coverage, e.g., inland seas such as the San Carlos waters, shown in the upper right subset panel. Some southern portions of West Falkland also recorded fewer instances of significant positive correlation, possibly because differences in coastline morphology, bathymetry, and wave exposure in West Falkland result in narrower, fringing beds closer to the coastline than in East Falkland. For example, nearshore canopy is more likely to be removed by our nearshore mask and may also be more sensitive to wave effects.

## Discussion

### Automated approaches for remote sensing of giant kelp

The physical and biological characteristics of FLK make this region well suited for satellite monitoring of giant kelp. For example, FLK kelp beds are relatively wide and extend further offshore than beds in many other regions. This is particularly noticeable on the eastern-facing coast, which receives less wave energy and features a broader, shallower shelf than the western-facing coast. The kelp beds in FLK are also dominated by a single species (i.e., giant kelp or *Macrocystis pyrifera*), although *Lessonia* spp. is also common within the understory or along the bed edges [15]. Finally, the absence of heavy industry in FLK, combined with low human population density, means that there are few sources of local pollution which can complicate atmospheric correction of satellite imagery through the generation of urban aerosols or the release of effluent into the coastal ocean.

The citizen science and automated spectral methods produced kelp classifications that were similar to expert classifications based on visual inspection (Fig. 3) and validation of kelp pixels within 1km coastline partitions (Fig. 4). Visual inspection revealed that the kelp bed classifications produced by citizen science consensus (FF8) had less granularity than the automated spectral classification (DTM), which more closely resembled the expert classifications in the shape of the classified kelp beds. The difference in granularity between the classification methods was anticipated because the citizen scientists produce shapefiles by drawing vectors around the kelp bed, a process that is more likely to produce rounded, less granular shapes compared to the pixel-by-pixel analysis of automated spectral methods.

Validation of the FF8 and DTM methods (Fig. 4) indicated better agreement of the DTM classifications with the expert dataset. Previously, Bell et al., 2020 [10] applied a correction to Landsat 8 (OLI) fractional coverage products to account for differences in spectral response between OLI and earlier sensors. The validation analysis herein did not indicate significant bias or difference in accuracy of kelp classifications between sensors, and applying a similar spectral response function resulted in over-prediction for the OLI products compared to ETM+ and TM. As a result, no spectral response function was applied to DTM for the OLI data products used herein. The difference in intersensor responses between the present study and Bell et al., 2020 [10] is most likely due to the conversion (in this study) to binary products. This would suggest that differences between sensors were less severe for low percent coverage values (i.e., those below the presence/absence threshold of 13%). The consistency between sensors could also be due, in part, to the spectral response correction already applied during USGS default processing, in which spectral reflectances are derived for central wavelengths.

Despite differences in accuracy between the citizen science and spectral automated methods, considering time-series using each of the FF8 and DTM data products provides useful redundancy. For example, the methods use different information from the satellite imagery, and therefore are susceptible to different causes of classification errors. The spectral automated method, which is derived pixel-by-pixel, does not incorporate spatial information (except by some filtering rules applied during post-processing), whereas the human viewers who produce the citizen science classifications recognize shape patterns, and therefore are significantly relying on spatial information. The spectral automated approach often produces false positives in individual pixels where the NIR domain is brightened (e.g., as caused by wave facets or suspended sediment). These errors are significantly mitigated by post-processing of the DTM data products, e.g., by removing pixels which rarely contain kelp or which are not significantly correlated with adjacent, kelp-containing pixels. As an alternate example, the citizen science approach falsely classified a large field of drifting debris as kelp canopy, despite a different spectral shape within the debris field.

The automated spectral method generated kelp classifications that were similar to the KD dataset [4], although with slightly better sensitivity, shown in Fig. 3. Differences in sensitivity are most likely due to the combination in DTM of the decision tree and the spectral unmixing methods. By pairing two approaches, the DTM dataset can use a more sensitive criteria (spectral unmixing detects fractional coverage) after rejecting pixels that are unlikely to contain kelp and therefore may generate false positives. Other reasons for the improved sensitivity of the DTM method include the increased number of spectral channels for the Landsat sensors compared to the Sentinel-2 sensor MSI, as well as the increased signal-to-noise characteristics for the former [42].

### Long-term variability in FLK giant kelp canopy

Recently, a global meta-analysis of regional kelp forest trends found that more kelp forests were decreasing in area than were increasing, but that regional variability was high [13]. Global observations of kelp forests are geographically and temporally patchy, with more research focusing on northern hemisphere regions (especially Southern Californian waters) despite evidence that kelp forest ecosystems within less studied regions of the southern hemisphere are anticipated to be similarly productive and highly dynamic [3]. For example, kelp forest declines in the Atlantic Ocean have primarily been reported from northern hemisphere sites, but there are insufficient observations from southern hemisphere regions to determine whether the southern Atlantic Ocean forests are more resilient than their northern Atlantic Ocean equivalents [14]. Improving the automation of kelp classification in satellite imagery (e.g., using the citizen science or automated spectral approaches tested herein) can expand the geographical extent of kelp canopy datasets and provides globally consistent observations dating back approximately three and a half decades (based on Landsat 5 imagery). These global observations support time-series analyses of previously under-studied regions in order to test whether these ecosystems have experienced sustained changes in canopy coverage. For example, regions like FLK in the southern Atlantic Ocean were not included in the Krumhansl et al., 2016 [13] global meta analysis due to a lack of kelp forest observations in those systems.

The timeseries presented herein using DTM (1985 to 2021) and FF8 (1997 to 2017) data products did not indicate significant trends in giant kelp canopy within the FLK region. Specifically, both the DTM and FF8 time series indicated slight (non-significant) declines, with P = 0.28 and P = 0.10, respectively. DTM canopy area binned within 1km coastline segments also did not produce cohesive spatial structure in the direction or magnitude of trends (Supplemental Fig. S3), and therefore the potential role of wave exposure or coastline orientation was not revealed by our time-series analyses. Stable giant kelp canopy area in the FLK region is consistent with recent work – in which imagery from satellites (2015-2020) as well as uncrewed (2019-2020) and crewed (1973) aircraft were compared with records from Charles Darwin and contemporaries (1829-1834) – that found that the locations of kelp beds recorded in the late 20^th^ and early 21^st^ centuries were in most instances consistent with those from 19^th^ century logs [15]. These results are also consistent with a recent report from the nearby Tierra del Fuego region of South America, in which present-day giant kelp density was found to be similar to that observed in a 1973 diver survey [43].

Although the timeseries did not provide evidence for a sustained decline in giant kelp canopy, the DTM dataset does capture a recent decrease in giant kelp canopy beginning approximately in 2017 and persisting through the end of the time series. The recent decrease leads to the lowest observed canopy area occurring during the final year of observations, and monitoring of this system should continue in order to detect potential further declines. However, the DTM dataset also indicates that a canopy area decline between 1985 and 1987 was similar in magnitude to the more recent decline, although there were many fewer observations during this earlier time period (see Supplementary Table S1), and observations in 1985 were only available from the western portion of the FLK region. The ability to identify historical precedent in the magnitude of the recent decline demonstrates the importance of obtaining long-term records when observing species like giant kelp with high interannual variability. For example, the DTM dataset included the 1987 low kelp observations and produced a higher P-value than the shorter FF8 dataset, which did not include observations before 1997. However, there was no historical precedent in either of our datasets for the lowest total canopy area recorded in 2021.

### Environmental drivers of giant kelp variability in FLK

Physical and biological factors that regulate the growth and survival of giant kelp include the temperature, nutrient content, and turbidity of the water, as well as the abundance of grazers [5]. Among these environmental drivers, temperature is routinely measured by ocean-viewing satellites at a variety of spatial and temporal scales. Grazer dynamics were not included in this study, which focused on continuous, spatially resolved datasets, but grazer population changes, e.g., of sea urchins that inhabit the FLK subtidal zone [34], have contributed to large kelp forest declines in other systems [44, 45].

Because of the latitude and oceanography of the FLK region, water temperatures are not anticipated to reach values associated with thermal stress in giant kelp, although local thermal adaptations can increase sensitivity to heat stress for individual populations [46]. We identified a positive correlation between synthetic nitrate and giant kelp canopy area, with the variability in giant kelp canopy lagging nitrate by one year. Overall, the correlations between nitrate and giant kelp canopy were not strongly dependent on coastline orientation, indicating that nitrate availability is important for giant kelp growth across the FLK region, and the effects were not severely diminished by variability in finer-scale, nearshore processes. Seasonal nitrate limitation has been reported for giant kelp beds within the FLK region [17], and a synthetic nitrate product derived from satellite SST was developed based on *in situ* relationships between temperature and nitrate measurements (Supplemental Fig. S2). Based on reports of nitrate limitation as well as low water temperatures, we present positive correlation of synthetic nitrate to lagged kelp canopy (P < 0.01 for the full FLK region) rather than negative correlation with temperature, although our nitrogen estimate is derived as a linear model of satellite SST.

Marine heatwave events (MHW) – high temperature episodes that coincide with lower nutrient availability and can facilitate the spreading of disease and non-native species [47] – have often preceded regional declines in canopy-forming kelp [44, 45, 48–50]. We did not detect a sustained trend or large annual anomalies in satellite estimates of nitrate or temperature during the period spanning 2002 to 2021 (SST data from the MODIS *Aqua* satellite began in 2002), which is consistent with our time-series analysis that did not detect changes in giant kelp canopy area. The one-year lag detected in the relationship between kelp canopy and synthetic nitrate suggests that adequate nutrient availability increases the health and reproductive success of individuals in order to seed the subsequent year’s population. The time lag is also consistent with the lag detected between ENSO variability and giant kelp in the Tierra del Fuego region [43].

Source waters to the FLK region are supplied primarily by the FC (Fig. 1), which transports cold ACC water northward. Stable oceanographic conditions in FLK during the observation period were likely due to the state of the FC, as well as, perhaps, the large distance separating the FLK region from the northward confluence of the FC with the warmer, more nutrient-poor BC waters [32]. For comparison with another region on a western subpolar ocean margin, a recent die-off of giant kelp in Tasmania coincided with a restructuring of oceanographic currents, which temporarily changed the source waters for the region such that nutrient availability decreased, temperatures increased, and an invasive species of sea urchin was introduced, which increased grazing on giant kelp fronds [44].

We did not detect an association between giant kelp canopy and ENSO, contrary to findings in the nearby Tierra del Fuego region in which kelp coverage was found to be negatively correlated to ENSO when lagged by one year [43]. Despite the proximity of eastern Tierra del Fuego and FLK (approximately 600km), this difference may be due to regional variation in current exposure. We also did not detect a significant association between kelp canopy and the Southern Annular Mode (SAM), despite the importance of this climate oscillation on atmosphere and ocean processes at latitudes spanning the FLK region. The SAM strongly influences temperatures and storm systems across the southern hemisphere, and long-term trends have been reported for the SAM during the second half of the 20^th^ century [51]. Although we did not detect a relationship between giant kelp canopy and the SAM, potential effects should be investigated further because of the coarse temporal resolution of the data products used in our analysis (e.g., quarterly or annual averages).

## Conclusion

Reports of kelp population declines have most often been associated with trailing range-edge populations, or those nearest the equator, because low latitude kelp forests are most likely to experience thermal stress and nutrient limitation related to climate warming [14]. However, mid-range populations are also susceptible to present and future warming trends [52], for example because of the introduction of invasive species that graze kelp [44], or because ecotypes can develop intraspecific thermal adaptations that increase sensitivity to temperature fluctuations [46]. Due to the complex effects of climate change, methods to improve the geographical coverage of kelp forest observations are needed in order to produce consistent, global-scale observations that include remote and understudied kelp forest environments.

Our study evaluated approaches for automating classification of giant kelp within Landsat imagery to enable more geographically expansive remote observations of kelp forests. We found that, despite differences in granularity, a citizen science consensus approach (FF8) and a spectral approach based on a decision tree paired with a spectral unmixing algorithm (DTM) each provided similar perspectives of kelp forest canopy variability in the FLK region. Based on differences in the types of information used in each approach (e.g., humans recognize spatial patterns, whereas DTM classifies individual pixels), future advancements that can incorporate spatial structure with a complete set of spectral information would be beneficial, e.g. computer vision.

We applied the automated methods to test for sustained changes in giant kelp within the FLK region and did not find evidence of long-term trends in canopy area using either approach. Our results were consistent with other recent work that included the FLK region [15], as well as work in the nearby Tierra del Fuego region [43], which also did not find evidence of long-term changes within these southern Atlantic Ocean kelp forest ecosystems. The regularity of satellite observations allowed for comparisons with ocean state variables, which revealed that strong associations between temperature or synthetic nitrate and giant kelp canopy lagged one year. Based on the region’s low maximum temperatures, these results suggest that nitrate variability is an important control of giant kelp canopy area in FLK.

Resources for *in situ* monitoring of coastal environments are less available for regions that are distant from major human population centers. Fewer observations are available from the south hemisphere in general, particularly from the southern Atlantic Ocean, despite the high productivity and the expansive area of giant kelp forests in these regions. Satellite imagery enables continuous and sustained observations of kelp forests at a global scale, but making use of these large datasets requires improvements in automation and image processing. Automated kelp classification tools – using citizen science annotations or a decision tree paired with a spectral unmixing algorithm – can provide accurate and routine observations of giant kelp canopy to test for environmental change in understudied coastal environments.

## Acknowledgments

Support for this work was provided by the National Aeronautics and Space Administration as part of the Citizen Science for Earth Systems Program (grant #80NSSC18M0103), and by the National Science Foundation through the Santa Barbara Coastal Long-Term Environmental Research program (grants OCE #0620276 & #1232779). We are extremely grateful for valuable feedback provided by (in alphabetical order): Paul Brewin (South Atlantic Environmental Research Institute), Neil Golding (Aquarius Survey and Mapping), and Michael Harte (Oregon State University).

## Supporting information

**S1.**
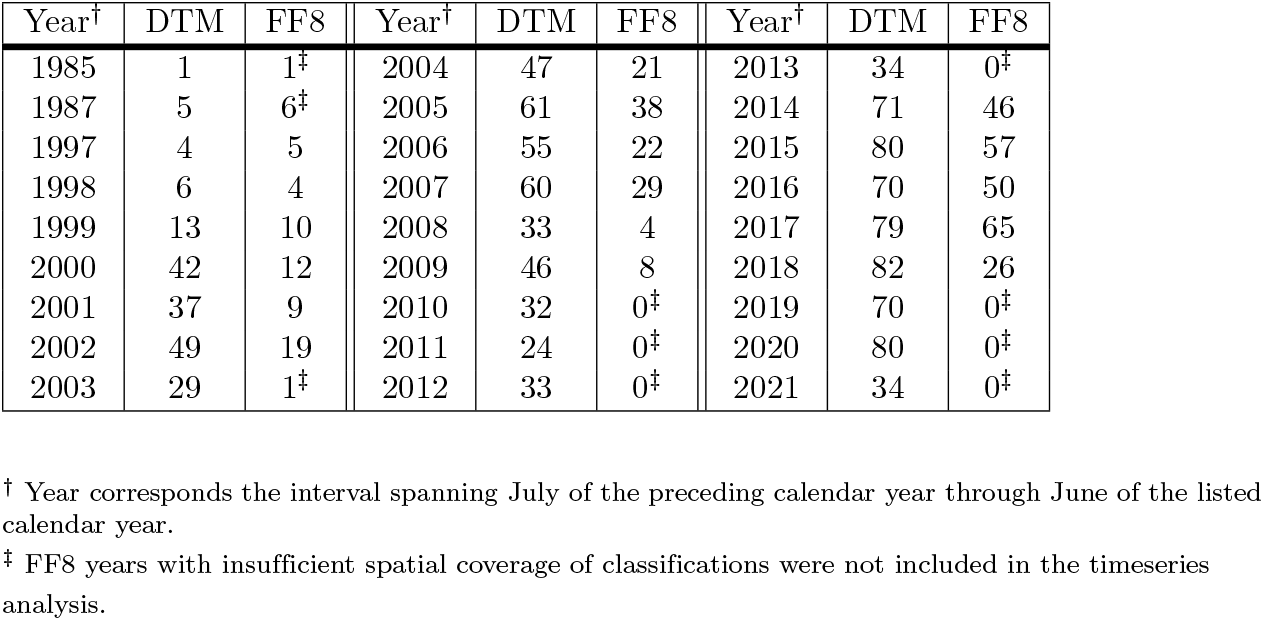
Annual observation days. Number of scenes observed during annual intervals (July-June) from the automated spectral (DTM) and citizen science (FF8) datasets.

**S2.**
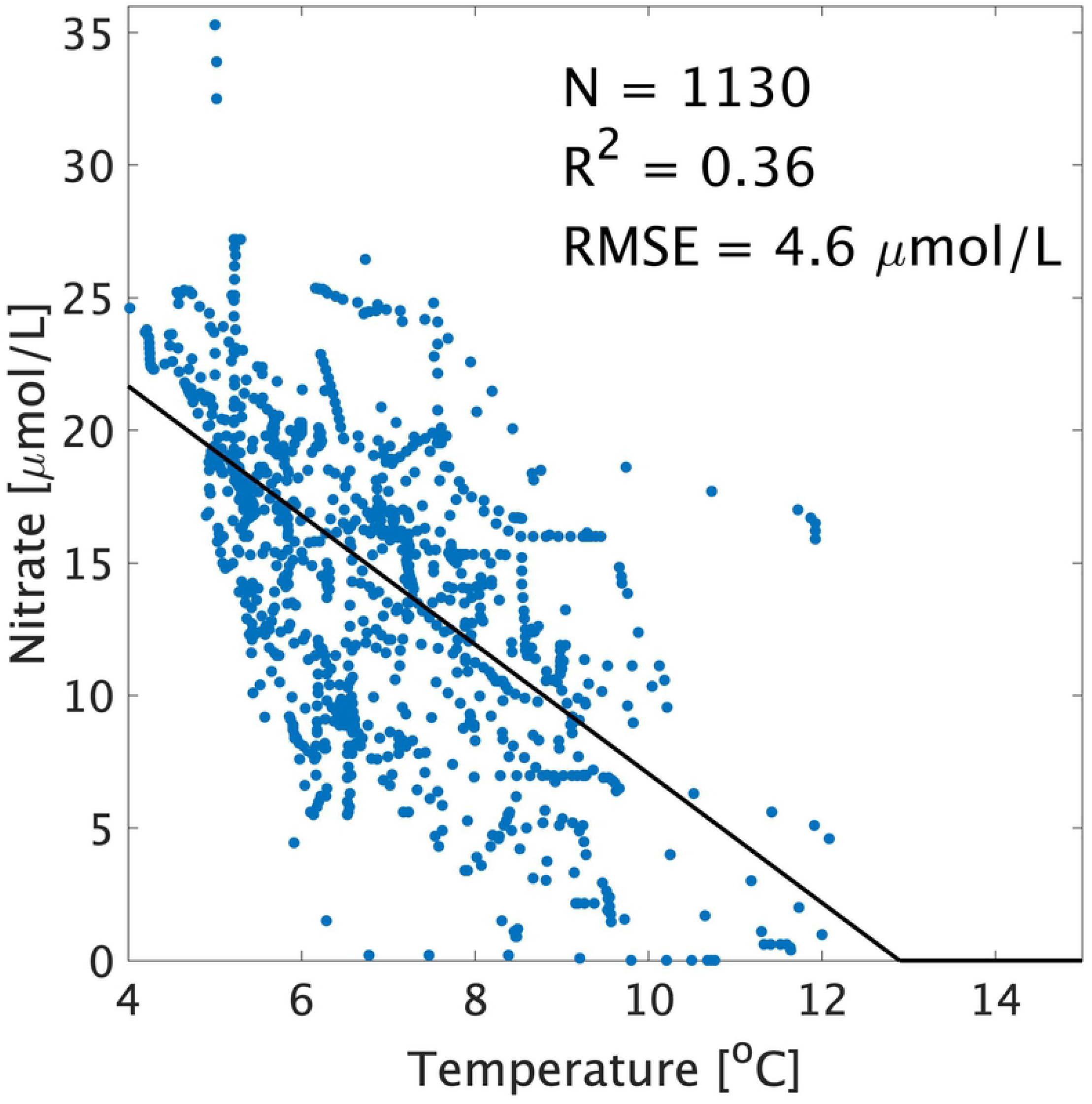
Derivation of synthetic nitrate. *In situ* nitrate and temperature measurements were obtained from the World Ocean Atlas within the region spanning 57.0° to 62.0° W and 50.0° to 53.0°. A synthetic nitrate model was derived from the *in situ* values using least-squares linear regression, and negative synthetic nitrate values were set to zero.

### Nitrate and temperature relationship for the FLK region

Nitrate and temperature measurements obtained from the World Ocean Atlas are shown as solid blue dots, and the synthetic nitrate model is overlaid as a solid black line.

**S3.**
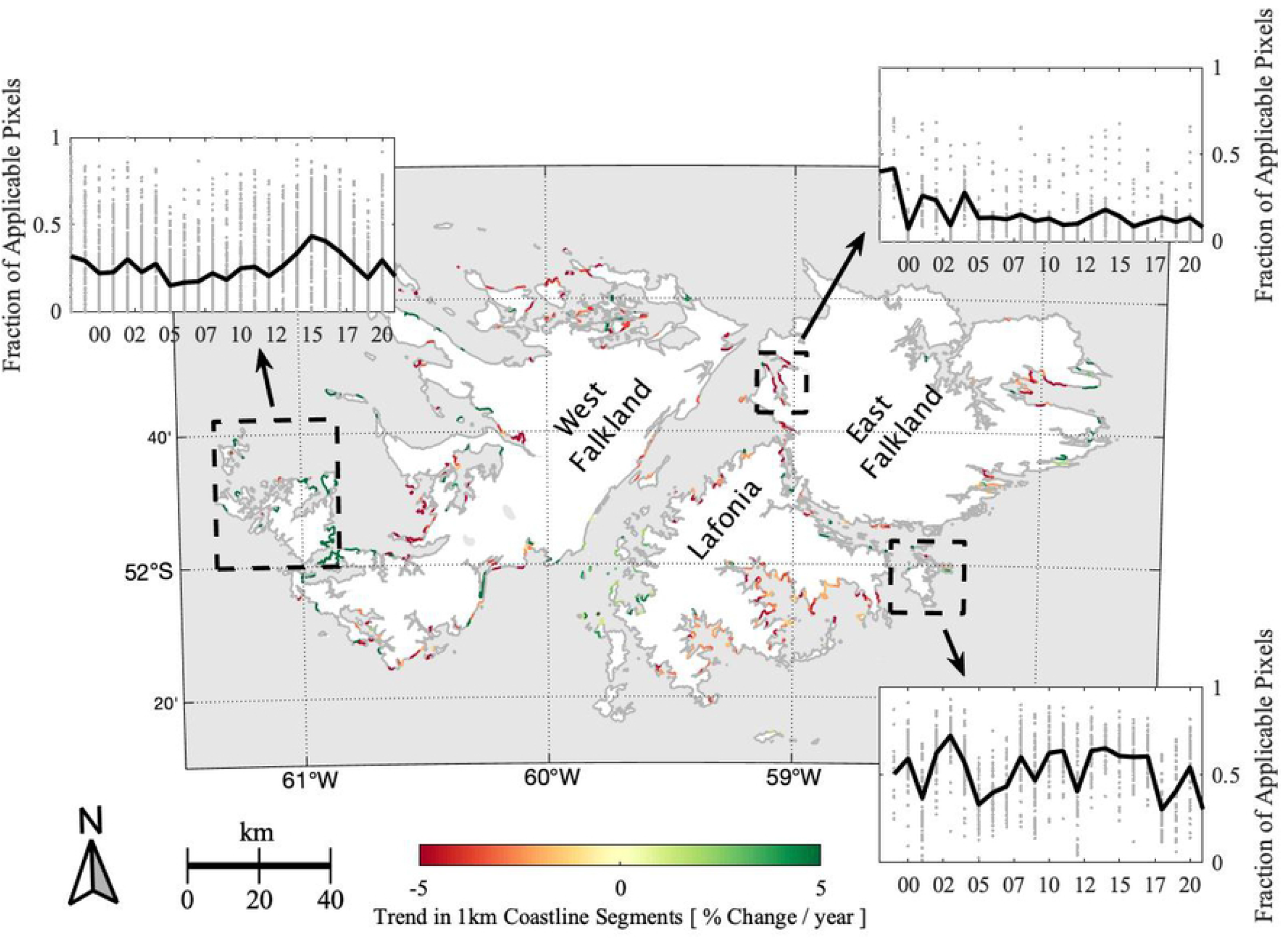
Regional trends in giant kelp canopy. Spatial variations in canopy trends were evaluated by aggregating annual DTM data products to their nearest 1km coastline segments and testing for long-term trends within the data products assigned to each segment.

### Trends in canopy extent partitioned to 1km coastline subsets

Significant (P < 0.01) linear trends in annual mean canopy coverage based on DTM data products are shown in green and red. Coastline subsets wherein trends are not significant (P ≥ 0.01) are indicated in gray. Timeseries of select coastline subsets are shown in the insets for regions indicated with a dashed black line, with year shown on the x-axis, as follows: Weddell Island and adjacent islands (upper left); San Carlos waters (upper right); and Lively Island (lower right).

